# Transcriptionally induced enhancers in the macrophage immune response to *Mycobacterium tuberculosis* infection

**DOI:** 10.1101/303552

**Authors:** Elena Denisenko, Reto Guler, Musa Mhlanga, Harukazu Suzuki, Frank Brombacher, Sebastian Schmeier

**Affiliations:** Massey University, Institute of Natural and Mathematical Sciences, Albany, Auckland, New Zealand.; Institute of Infectious Diseases and Molecular Medicine (IDM), Division of Immunology and South African Medical Research Council (SAMRC) Immunology of Infectious Diseases, Faculty of Health Sciences, University of Cape Town, Cape Town, South Africa.; International Centre for Genetic Engineering and Biotechnology (ICGEB), Cape Town Component, Cape Town, South Africa.; Gene Expression & Biophysics Group, CSIR Synthetic Biology ERA, Pretoria, South Africa.; Institute of Infectious Diseases & Molecular Medicine (IDM), Division of Chemical Systems & Synthetic Biology, University of Cape Town, Cape Town, South Africa.; Gene Expression and Biophysics Unit, Instituto de Medicina Molecular, Faculdade de Medicina Universidade de Lisboa, Lisbon, Portugal.; Division of Genomic Technologies, RIKEN Center for Integrative Medical Sciences, 1-7-22 Suehiro-cho, Tsurumi-ku, Yokohama, Japan.

**Keywords:** eRNA, macrophages, transcriptional enhancers, transcriptional regulation, tuberculosis

## Abstract

**Background:** Tuberculosis is a life-threatening infectious disease caused by *Mycobacterium tuberculosis* (*M.tb*). *M.tb* subverts host immune responses to build a favourable niche and survive inside of host macrophages. Macrophages can control or eliminate the infection, if appropriate transcriptional programs are activated. The role of transcriptional enhancers in the activation and maintenance of these programs remains unexplored.

**Results:** We analysed transcribed enhancers in *M.tb*-infected mouse bone marrow-derived macrophages. We established a link between known *M.tb*-responsive transcription factors and transcriptional activation of enhancers and their target genes. Our data suggest that enhancers might drive the macrophage response via transcriptional activation of key immune genes, such as Tnf, Tnfrsf1b, Irg1, Hilpda, Ccl3, and Ccl4. We report enhancers acquiring transcription *de novo* upon infection. Finally, we link highly transcriptionally induced enhancers to the activation of genes with previously unappreciated roles in *M.tb* infection, such as Fbxl3, Tapt1, Edn1, and Hivep1.

**Conclusions:** Our findings extend current knowledge of the regulation of macrophage responses to *M.tb* infection and provide a basis for future functional studies on enhancer-gene interactions in this process.

## Background

Tuberculosis (TB) remains a significant global threat, which causes over one million deaths each year. The causative agent of TB is *Mycobacterium tuberculosis* (*M.tb*), an intracellular pathogen that mainly persists inside host macrophages [1]. Over 30% of the world’s population is infected with *M.tb*, and the infection progresses to active TB in about 5-10% of cases [1, 2]. Macrophages are one of the first lines of a host’s defence against invading bacterial pathogens [3]. The complex interplay between host macrophages and *M.tb* is believed to be central to the control of infection and defines the infection outcome [4, 5]. Macrophages are equipped with a multitude of strategies to combat *M.tb*, however, the pathogen has developed a wide range of matching resistance mechanisms, allowing it to avoid destruction and to survive and proliferate inside macrophages [5]. Hence, macrophage responses need to be tightly controlled in order to eliminate the pathogen. The lack of effective TB control systems is in part explained by significant gaps in our knowledge of the biology of *M.tb* and its interactions with the host [4]. Consequently, understanding the cellular pathways that underlie the initial infection and TB progression remains a scientific challenge directly applicable to human health.

Gene expression in eukaryotic cells is a complex process guided by a multitude of mechanisms [6]. Regulation of transcription represents one of the first layers of gene expression control, which largely defines rapid signal-dependent expression changes [7]. Enhancers are defined as *cis*-regulatory DNA regions that activate transcription of target genes in a distance- and orientation-independent manner [8]. Nowadays, enhancers are considered major determinants of gene expression programmes required for establishing cell-type specificity and mediating responses to extracellular signals [9-11].

Enhancers are characterised by a set of distinctive features. Genomic regions surrounding enhancers carry a combination of H3K4me1 and H3K27ac histone marks that has been considered an enhancer-specific chromatin signature [12, 13]. H3K4me1 demarcates established or primed enhancers, which may or may not be active, while a combination of H3K4me1 and H3K27ac marks active enhancers [12, 13]. Enhancer regions carry multiple DNA binding sites and can recruit transcription factors and coactivators, RNA polymerase II and other proteins, such as histone acetyltransferases [9, 14, 15]. Enhancers serve as a platform for assembly of the transcription pre-initiation complex, which can result in enhancer regions being transcribed into non-coding enhancer RNAs termed eRNAs [14, 15]. This novel class of RNAs was first introduced in a genome-wide study in mouse neurons [16]. Later on, a number of studies showed that the production of eRNAs correlated with target mRNA synthesis and eRNAs could serve as robust and independent indicators of active enhancers, that are more likely to be validated *in vitro* [17-21]. Detectable eRNA levels are usually low, possibly due to their short half-life and fast degradation by RNA exosomes or their generally low transcription initiation rates [11, 22-24]. Nevertheless, eRNA transcription can be used for a genome-wide identification of active enhancers [17, 25, 26].

The dominant model of transcriptional regulation by enhancers states that it is exerted via direct physical interaction between an enhancer and a target gene promoter, mediated by DNA looping [8]. Topologically associating domains (TADs) have emerged as critical conserved units of chromatin organisation that favour internal DNA contacts, whereas regulatory interactions between TADs are limited [27, 28]. Enhancer-promoter contacts are believed to occur almost exclusively within the well-conserved TADs [29]. Notably, enhancer-promoter interactions are not limited to one-to-one contacts. Instead, an enhancer might regulate a few genes, and multiple enhancers might contribute to the activation of a gene [30]. Such enhancer redundancy was recently shown to confer phenotypic robustness to loss-of-function mutations in individual enhancers [31]. Both enhancers and enhancer-gene regulatory interactions are characterised by a remarkable tissue specificity [13, 18, 20]. Such tissue specificity is crucial for establishing cell-type- and state-specific transcriptional programmes [9, 10]. Moreover, enhancer-gene interactions can be dynamically rewired in response to environmental stimuli, enabling fine tuning of gene expression programmes [19, 32].

Previously we used cap analysis of gene expression (CAGE) and epigenetic data to identify on large-scale transcribed enhancers (i.e. enhancers producing eRNAs) in bone marrow-derived mouse macrophages (BMDM) [33]. We have established a transcribed enhancer and target gene interactome and characterised the roles of enhancers in guiding macrophage polarisation into distinct pro- and anti-inflammatory phenotypes [33]. Here, we extended the former study to conduct the first to our knowledge genome-wide analysis of transcribed enhancers guiding BMDM response to *M.tb* infection. Our findings indicate that transcribed enhancers are extensively involved in the induction of immune genes during *M.tb* infection. We identify and characterise enhancers with induced or *de novo* acquired eRNA expression and transcription factors that likely drive these changes. We report enhancer regions that target known immune genes crucial for the host response to *M.tb*. These findings are extended by highlighting genes with previously unappreciated roles in *M.tb* infection, as their regulation by many enhancers points to their functional importance. Taken together, our findings extend the current knowledge of *M.tb*-induced immune response regulation in macrophages and provide a basis for future functional studies on enhancer-gene interactions in this process.

## Results

### Transcribed enhancers in macrophage responses to *M.tb* infection

We analysed the host transcriptional response to *M.tb* infection in mouse bone marrow-derived macrophages (BMDM) at 4, 12, 24, and 48 hours post infection (see Methods). Non-infected control BMDM were profiled prior to infection (0 h) and at matched time points (4, 12, 24 and 48 h). First, we analysed overall gene expression changes and found that they were the strongest at 4 h post infection and declined with time (Fig 1a-c). Half as many differentially expressed genes (DEGs) were detected at 12 h as at 4 h, and almost no genes were significantly differentially expressed at 24 or 48 h post infection (see Methods, Fig 1a). We combined the DEGs from all time points into two unique lists of 1,384 up- and 1,604 down-regulated DEGs for further analysis.

**Fig 1.**
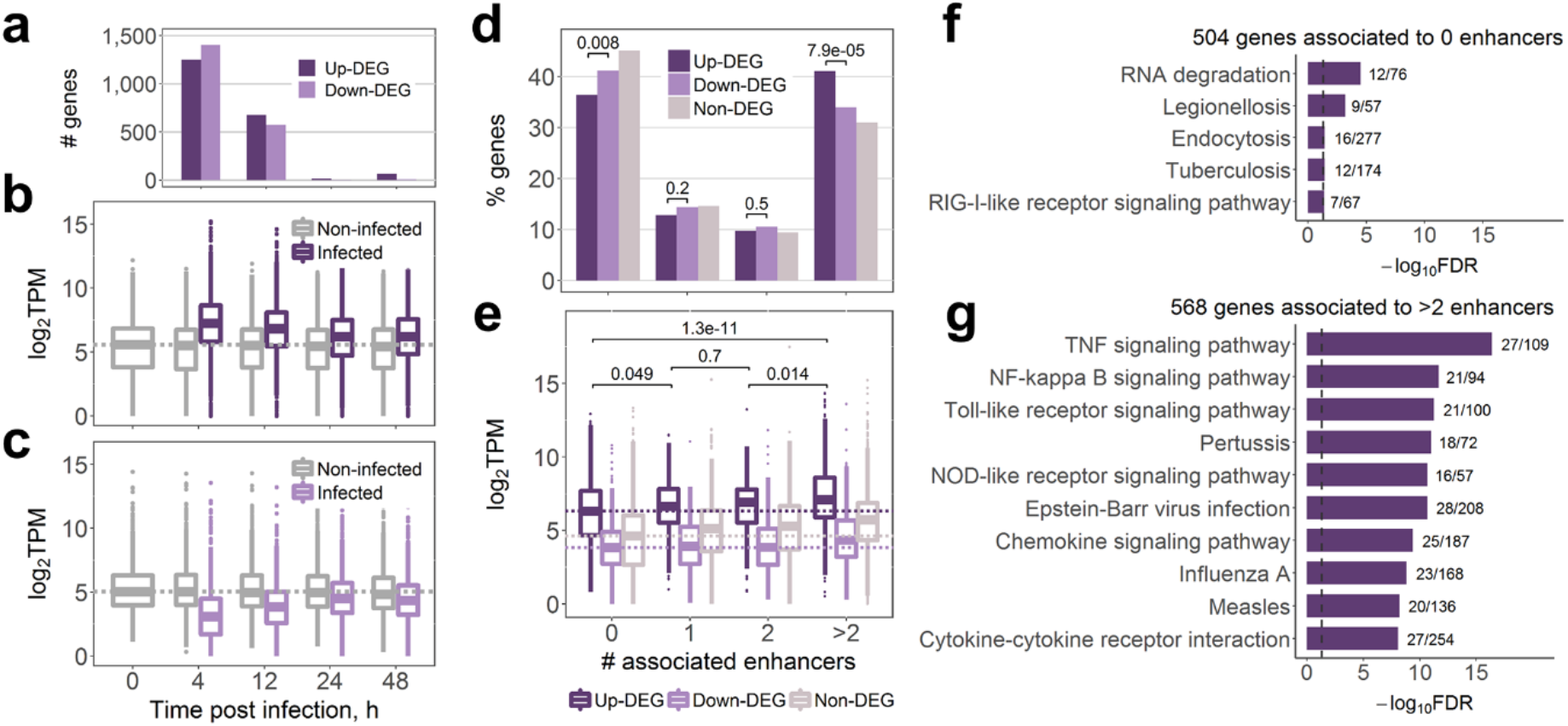
Enhancers mediate up-regulation of immune genes in macrophages upon *M.tb* infection. (a) Numbers of differentially expressed genes (DEGs) in infected macrophages vs. macrophages prior to the infection (0 h). (b) Expression of 1,384 up-regulated DEGs. (c) Expression of 1,604 down-regulated DEGs. In (b) and (c), genes are differentially expressed at any time point; expression in TPM was averaged across replicates; dashed lines show median gene expression prior to the infection. (d) Percentage of genes associated with different number of enhancers in infected macrophages; numbers indicate Fisher’s exact test p-values. (e) Expression of genes associated with different number of enhancers in infected macrophages; expression in TPM was averaged across infected samples, dashed lines show the median expression of genes not associated with any transcribed enhancer; numbers indicate Wilcoxon two-sided rank sum test p-values. (f) KEGG pathway maps significantly enriched for up-regulated DEGs with no associated transcribed enhancers, FDR < 0.05. (g) Top 10 KEGG pathway maps with the lowest FDR enriched for up-regulated DEGs associated with more than two transcribed enhancers. In (f) and (g), next to the bars are the numbers of genes in the KEGG term covered by our gene list; dashed lines indicate FDR = 0.05.

We have previously identified 8,667 actively transcribed enhancers and their target genes in mouse BMDM [33]. Here, we found that many of these enhancers acquired higher eRNA expression in response to *M.tb* infection (S1a Fig). Moreover, enhancers associated with up-regulated DEGs in infected macrophages showed an increase in eRNA expression (S1b Fig, see Methods and S1 Table for the list of up-regulated DEGs and their enhancers). Hence, BMDM enhancers showed an overall increase in transcriptional activity upon *M.tb* infection.

We investigated the differences in the enhancer repertoire between DEGs and non-DEGs to uncover the role of enhancers in the *M.tb* infection response. Genes with no transcribed enhancers composed 36.4% of up-regulated DEGs, whereas this percentage was significantly higher at 41.1% for down-regulated DEGs (Fisher’s exact test two-sided p-value 0.008) (Fig 1d). Furthermore, 41% of up-regulated DEGs, but only 34% of down-regulated DEGs were associated with more than two transcribed enhancers (Fisher’s exact test two-sided p-value 7.9*10^-05^) (Fig 1d). Finally, non-DEGs had the highest percentage of genes with no transcribed enhancers (45%) and the lowest percentage of genes with more than two enhancers (31%) (Fig 1d). Hence, transcribed enhancers likely play a prominent role in up-regulation of protein-coding genes in the response to *M.tb* infection.

Previously we have shown that regulation of genes by many transcribed enhancers in BMDM was a concomitant of higher gene expression and tissue-specific function [33]. Here, we asked whether the same properties could be observed for up-regulated DEGs, as genes most likely to be involved in the elimination of *M.tb*. Indeed, as before, we noted higher expression levels in genes associated with more enhancers in *M.tb*-infected macrophages (Fig 1e). Gene set enrichment analysis (GSEA, see Methods) showed that DEGs with no transcribed enhancers in *M.tb*-infected macrophages were only significantly enriched (FDR < 0.05) in five KEGG pathway maps (Fig 1f). In contrast, genes associated with more than two enhancers were significantly enriched in as many as 92 pathway maps (S2 Table), and showed a much stronger enrichment for more specific infection-related pathways (Fig 1g, S2 Table) when compared to genes with no enhancers (Fig 1f). The enrichment analysis points to the assumption that up-regulated DEGs without transcribed enhancers are functionally less related than those associated with more than two actively transcribed enhancers. Moreover, these results indicate that even within such a process-oriented set as the list of up-regulated DEGs, multiple enhancers might regulate the most highly expressed and functionally important genes. We repeated this analysis for all genes (as opposed to only DEGs) and their associated enhancers in infected macrophages and observed a similar trend (S2 Fig), in agreement with our previous study [33].

We next compared our transcribed enhancers to a set of inflammation-sensitive LPS-responsive macrophage super enhancers (SEs) reported by Hah et al. [34]. Super-enhancers (or stretch enhancers) have emerged as a sub-class of particularly potent enhancers, which are associated with higher levels of enhancer-specific histone marks and regulate key cell identity genes [35, 36]. Among 2,999 enhancers associated with up-regulated DEGs, 45.9% overlapped SE regions. This percentage was significantly lower at 30% for the remainder of our BMDM transcribed enhancers [33] (two-sided Fisher’s exact test p-value < 2.2*10^-16^, odds ratio 1.98). Interestingly, of 880 up-regulated DEG associated with transcribed enhancers, 477 were associated with enhancers overlapping SEs, and these DEGs showed a much stronger enrichment for immune-related functions, when compared to the 403 DEGs for which none of their associated enhancers overlapped SEs (S3 Fig).

Taken together, our findings indicate that the up-regulation of immune genes in BMDM upon *M.tb* infection might be largely driven by transcribed enhancers. Comparison of the three subsets of up-regulated DEGs showed the strongest enrichment for specific immune response pathways in up-regulated DEGs associated with SEs (S3b Fig) and the weakest enrichment in up-regulated DEGs not associated with any transcribed enhancers (Fig 1f), highlighting the functional importance of SEs in BMDM response to *M.tb* infection.

### Transcriptionally induced enhancer regulation of immune genes during *M.tb* infection

We further set out to investigate a subset of enhancers that targeted up-regulated DEGs and were themselves highly transcriptionally induced upon infection. We focused on 809 DEGs that were associated to transcribed enhancers and up-regulated at 4 h post infection, as we observed the strongest transcriptional response upon infection at this time point. Of enhancers targeting these DEGs, we selected those with the highest eRNA expression at 4 h and its fold change as compared to 0 h, by requiring both these values to be in the upper quartiles of their corresponding distributions (see Methods). The derived set of 257 enhancers (further referred to as induced enhancers) was associated with 263 of 809 DEGs that were up-regulated at 4 h and associated with transcribed enhancers (S4 Fig, S3 Table). We investigated expression of the induced enhancers in other mouse tissues (S4 Table). Interestingly, we found that the set of enhancers showed the highest average and maximum eRNA expression, as well as the highest percentage of samples with nonzero eRNA expression in infected macrophages (S5 Fig). In addition, induced enhancers were over-represented in SE regions [34] when compared to the remainder of BMDM enhancers, with 60.7% of the induced enhancers overlapping SEs as compared to 34.7% of non-induced enhancers (two-sided Fisher’s exact test p-value < 2.2*10^-16^, odds ratio 2.9). These findings indicate a high specificity of the induced enhancers to the BMDM infection response and highlight the fact that they are likely key elements for driving the transcriptional responses of the macrophage upon infection.

Next, we investigated DEGs that were targeted by many induced enhancers as it stands to reason that these genes play crucial parts in the response to *M.tb*. Among the 263 DEGs, Tumour necrosis factor receptor 2 (*Tnfrsf1b*) was associated with the highest number of the induced enhancers, eight (Fig 2). Interestingly, one of the induced enhancers (chr4:145245568..145245969, Fig 2b) showed the second highest mean eRNA expression (28.79 TPM) at 4 h post infection among all enhancers targeting up-regulated DEGs. Tumour necrosis factor (*Tnf*), coding a ligand of Tnfrsf1b, was associated with three induced enhancers with mean eRNA expression of 2.4, 3.9, and 10.8 TPM at 4 h post infection. We found that induced enhancers associated with *Tnfrsf1b* were significantly over-represented in the corresponding TAD (eight induced enhancers among 38 BMDM enhancers in the TAD, hypergeometric test FDR = 0.005, see Methods). Interestingly, *Tnfrsf1b* was the only up-regulated DEG within the TAD (log_2_FC = 2.2 at 4 h vs. 0 h, Fig 2a) and encodes the Tnf receptor 2, which is known to interfere with apoptosis [37] and sensitize macrophages for Tnfr1-mediated necroptosis, a programmed form of inflammatory cell death resulting from cellular damage or infiltration by pathogens [38]. Given that all of *Tnfrsf1b*’s induced enhancers coincide with a SE, we hypothesise that the activation of the SE upon infection is driving the process in conjunction with increased eRNA expression from the induced enhancers.

**Fig 2.**
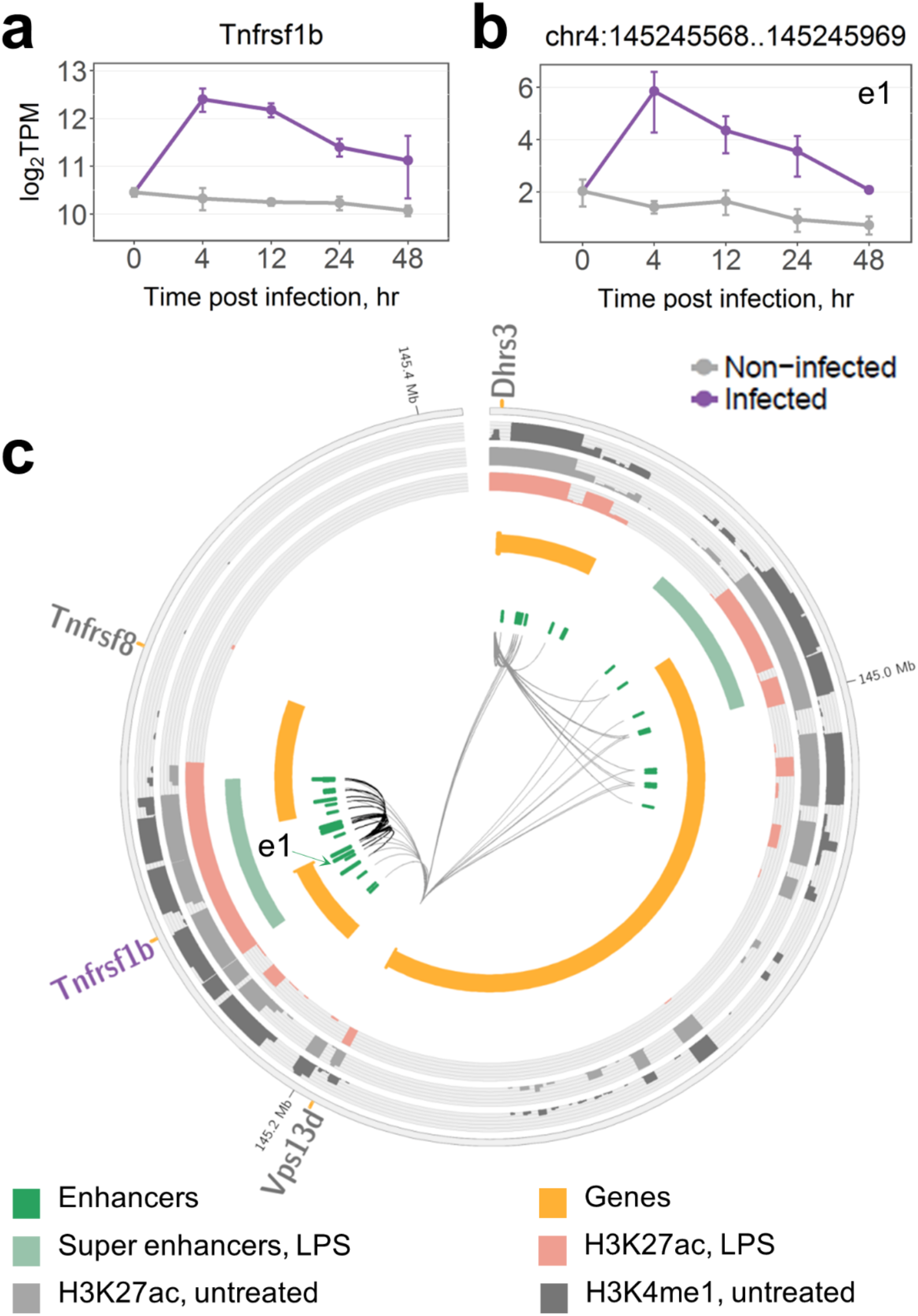
Regulation of Tnfrsf1b by induced enhancers. (a) Time course expression of the *Tnfrsf1b* gene. (b) Time course eRNA expression of *Tnfrsf1b*-associated induced enhancer. In (a) and (b), data were averaged over replicates and log-transformed, error bars are the SEM (see Methods). (c) TAD containing *Tnfrsf1b* and associated enhancers; induced enhancers are shown as longer green blocks. Genes are split into two tracks based on the strand, wide orange marks denote gene promoters. DEGs significantly up-regulated at 4 h are shown in purple and their associations with enhancers are shown as thicker black connections. Super enhancers are shown as defined by Hah et al. [34] in LPS-treated macrophages. Histone marks are shown as defined by Ostuni et al. [61] in untreated and LPS-treated macrophages.

Another TAD on chromosome 14 contained a group of three co-regulated DEGs (*Irg1*, *Cln5*, and *Fbxl3*) associated with six induced enhancers each, the second highest number after *Tnfrsf1b* reported above (S6 Fig). Moreover, among these six, the chr14:103037012..103037413 enhancer showed the highest mean eRNA expression (36.68 TPM) at 4 h post infection among all enhancers of up-regulated DEGs (S6b Fig, enhancer e2). Finally, six out of 14 enhancers in the TAD were deemed induced enhancers (significant over-representation with hypergeometric test FDR = 0.001, see Methods). Of the three DEGs, *Irg1* showed the strongest induction of log_2_FC = 5.2 at 4 h vs. 0 h (S6a Fig). Irg1 was recently shown to link cellular metabolism with immune defence by catalysing the production of itaconic acid, which has antimicrobial activity and inhibits the growth of *M.tb* [39]. Another gene in this TAD encodes Cln5 (log_2_FC = 2), which is required to recruit and activate Rab7 [40], a GTPase essential for phagosome maturation, a process which is crucial for microbial killing by macrophages and which can be disrupted by *M.tb* as a part of its survival strategy [41-43]. The link between highly induced enhancers and *Irg1* and *Cln5* points to biological processes important for the host response that might be driven by transcribed enhancers, while the immune functions of *Fbxl3* (log_2_FC = 1.4) are yet to be elucidated.

Induced enhancers were significantly over-represented with FDR < 0.05 in four more TADs, which we further investigated as potentially important *M.tb*-responsive genomic regions (S5 Table). One of the TADs (FDR = 0.001, five induced enhancers among eight BMDM transcribed enhancers, S7 Fig) is as large as 1.2 Mb and contains multiple genes, however, only *Hilpda* (*Hig2*) was differentially expressed and up-regulated at 4 h (log_2_FC = 6, S7a Fig). Hilpda is induced in hypoxia and is crucial to lipid accumulation in macrophages [44], which provides a favourable environment for dormant *M.tb* and might, thus, contribute to *M.tb* survival within the host [45]. Similarly, *Itgb8* was the only up-regulated DEG (log_2_FC = 7.1) in another TAD with five induced enhancers among 14 BMDM transcribed enhancers (FDR 0.011, S8 Fig). Although specific roles of Itgb8 in *M.tb* infection response have not yet been established, integrin alpha(v)beta8 is known to activate TGF-beta [46], an important mediator of susceptibility to *M.tb* [47].

A TAD with four induced enhancers among eight BMDM transcribed enhancers (FDR = 0.012) contains three DEGs up-regulated at 4 h post infection (S9 Fig). Cd38 and Bst1 (Cd157) are homologous NAD(+) metabolic enzymes up-regulated by Tnf [48], and Cd38 was shown to be involved in phagocytosis [49] and response to intracellular pathogen *Listeria monocytogenes* [50] in mouse macrophages. The role of the third gene in that TAD, transmembrane protein Tapt1, remains to be elucidated.

Finally, a TAD with five induced enhancers among 17 BMDM transcribed enhancers (FDR = 0.02) covers four DEGs *Ccl3*, *Ccl4*, *Ccl9*, and *Wfdc17* (S10 Fig). Ccl3 and Ccl4 are macrophage-derived inflammatory chemokines that induce chemotactic mobilization of immune cells [51], while Wfdc17 might have the opposite function decreasing production of pro-inflammatory cytokines [52], and the function of Ccl9 in macrophage infection response remains to be uncovered [51].

Taken together, these examples highlight six TADs (S5 Table), located on six different chromosomes, which show strong responses to *M.tb* infection and contain genes with both known and previously unappreciated roles in *M.tb* infection. These genes are under the control of multiple *M.tb* induced enhancers, which might be essential for contributing to the genes’ activation states.

To get further insights into the capacity of induced enhancer regulation during the response to *M.tb* infection, we investigated target DEGs of induced enhancers that were significantly enriched in particular biological pathways (S4b Fig). The Tnf signalling pathway showed the strongest enrichment for induced enhancer-regulated DEGs and included 18 DEGs up-regulated at 4 h and associated with the induced enhancers. Among these genes, in addition to *Tnfrsf1b* reported above, we identified *Tnf* itself, Tnf signalling pathway mediator *Traf5* and multiple effector genes targeted by induced enhancers (S6 Table). Tnf-alpha receptors are known to trigger the NF-kB signalling pathway, which was also enriched for DEGs regulated by induced enhancers, including receptors *Cd14* and *Cd40*, ligand *Il1b*, and TFs of canonical NF-kB signalling, *Nfkb1* and *Rela* (S6 Table). ‘Tuberculosis’ KEGG pathway map comprised five signal transduction mediators, *Irak2*, *Jak2*, *Malt1*, *Ripk2*, and *Src*, regulated by induced enhancers (S6 Table). In addition, induced enhancers target the *Eea1* gene, which is known to be involved in phagosome maturation, a process necessary for killing of bacteria within phagosomes [53] (S6 Table). Notably, genes encoding negative regulators of the listed signalling pathways, *Nfkbia*, *Tnfaip3*, and *Socs3*, were also associated with one to five induced enhancers (S6 Table), and showed up-regulation.

### Transcriptionally induced enhancers are enriched for immune transcription factor binding sites

Transcription factor (TF) binding motif analysis was performed to uncover TFs potentially involved in the transcriptional activation of induced enhancers. We identified twelve significantly over-represented motifs of TFs that were differentially expressed and up-regulated at 4 h post infection (see Methods, Table 1). Five of these motifs belong to the AP-1 family of TFs, among which the highest expressed one was *Junb*, recently reported to be an important regulator of immune genes in macrophages treated with LPS [54]. Interestingly, a negative regulator of AP-1, Jdp2, was also among the significantly over-represented motifs, although it was found only in 20.6% of the induced enhancers. Three motifs of NF-kB family were identified, among which *Rela* was reported above to be itself regulated by the induced enhancers, potentially forming a positive feedback loop. For another TF identified here, Irf1, we have previously reported that in association with Batf2 (log_2_FC = 2.7) it induced inflammatory responses in *M.tb* infection [55]. Both AP-1 and NF-kB families of TFs, as well as Irf1, play important roles in macrophages and can be triggered by a range of infection response receptors including Toll-like and Nod-like receptors [56, 57]. Rbpj, which showed the second strongest motif over-representation, is a key TF of canonical Notch signalling pathway, which is known to be activated by Toll-like receptor signalling pathways [58]. Finally, Nfe2l2 (Nrf2) regulates cytoprotective genes that enhance cell survival and was shown to increase phagocytic ability of macrophages and to improve antibacterial defence [59, 60].

**Table 1.** TF motifs over-represented in the induced enhancers.

| TF Motif | # overlapping enhancers | Expression, TPM | $\log_2FC$ | FDR |
| --- | --- | --- | --- | --- |
| FOSL1::JUNB | 118 (45.9%) | 19.4 / 523.7 | 5.4 / 2.6 | 2.7e-03 / 1e-04 |
| RBPJ | 117 (45.5%) | 295.7 | 1.8 | 7.3e-03 |
| REL | 96 (37.4%) | 165.3 | 3 | 4.3e-06 |
| FOSL2::JUNB | 91 (35.4%) | 81.2 / 523.7 | 2.4 / 2.6 | 1.2e-06 / 1e-04 |
| IRF1 | 88 (34.2%) | 1099.8 | 2.8 | 1.5e-04 |
| RELA | 87 (33.9%) | 309 | 1.7 | 1e-06 |
| JUNB | 84 (32.7%) | 523.7 | 2.6 | 1e-04 |
| FOSL1 | 80 (31.1%) | 19.4 | 5.4 | 2.7e-03 |
| FOSL2 | 80 (31.1%) | 81.2 | 2.4 | 1.2e-06 |
| Nfe2l2 | 54 (21%) | 684 | 1.5 | 2.4e-03 |
| JDP2 | 53 (20.6%) | 67.7 | 2.9 | 4.8e-04 |
| NFKB2 | 28 (10.9%) | 496.3 | 2.7 | 2.5e-04 |

Columns show TF motif name, number and percentage of overlapping enhancers among the induced enhancers, average expression of the corresponding TF(s) in infected BMDM at 4 h, fold change and FDR of differentially expressed test for the corresponding TF(s) in infected BMDM at 4 h versus non-infected control at 0 h. Motifs were retained for TFs with significant up-regulation at 4 h.

Importantly, 89.1% of the 257 induced enhancers considered here carry at least one of the twelve motifs, and these enhancers target 95.1% of the 263 up-regulated DEGs (Table 2). Among the motifs, AP-1 family members covered the largest percentages of the induced enhancers and their target genes, followed by the NF-kB family and Rbpj TF, highlighting their importance in enhancer regulation of *M.tb* response. We compared this TF regulation of protein-coding genes via enhancers to TFs that bind directly to the promoters of the 263 up-regulated DEGs (see Methods). In the promoters, Irf1, as well as AP-1, and NF-kB families were similarly significantly over-represented, whereas, Rbpj, Nfe2l2 and Jdp2 were not deemed significant and, thus, might be specific to the transcriptionally induced enhancers. Taken together, these findings link *M.tb*-perturbed signalling pathways and their key TFs to transcriptional activation of the induced enhancers, which in turn activate their immune target DEGs.

**Table 2.** TF-mediated regulation of genes via induced enhancers.

| TF Motifs | # overlapping enhancers | # target DEGs |
| --- | --- | --- |
| AP-1 (FOSL1::JUNB, FOSL2::JUNB, JUNB, FOSL2, FOSL1) | 128 (49.8%) | 180 (68.4%) |
| NF-κB (REL, RELA, NFKB2) | 117 (45.5%) | 157 (59.7%) |
| RBPJ | 117 (45.5%) | 160 (60.8%) |
| IRF1 | 88 (34.2%) | 126 (47.9%) |
| Nfe2l2 | 54 (21%) | 86 (32.7%) |
| JDP2 | 53 (20.6%) | 92 (35%) |
| Total (12 motifs) | 229 (89.1%) | 250 (95.1%) |

Columns show individual TF motifs or their groups, number and percentage of overlapping enhancers among the induced enhancers, number and percentage of DEGs targeted by these enhancers among the 263 DEGs up-regulated at 4 h and associated with the induced enhancers.

### A subset of enhancers is transcribed *de novo* upon *M.tb* infection

Interestingly, among 257 induced enhancers we found 17 enhancers that showed zero eRNA expression in all of the 22 non-infected macrophage samples. Hence, transcription of these enhancers was specifically acquired *de novo* in macrophages upon *M.tb* infection. These enhancers were associated with 31 of the 263 DEGs under investigation, which included *Hilpda*, *Il1b*, *Itgb8*, *Jak2*, *Src*, and *Tnfaip3* genes, reported above. We set out to further investigate in more detail the phenomenon of *de novo* transcription at enhancers.

We focused on enhancers that were transcriptionally silent in naïve BMDM, but acquired transcriptional activity *de novo* in *M.tb*-infected macrophages (further referred to as acquired enhancers). We hypothesized that such enhancers might either loop towards their target promoters in non-infected macrophages without being transcriptionally active, or form a novel DNA loop upon infection (Fig 3a-b). In total, we identified 356 acquired enhancers (see Methods). Their eRNA expression was the highest at 4 and 12 h post infection and declined with time (Fig 3c, left panel), in agreement with the DEG expression reported above. Notably, overall expression of acquired enhancers in infected macrophages was lower than that of induced enhancers (median of 0.23 TPM versus 1.73 TPM at 4 h). However, similarly to induced enhancers, acquired enhancers showed the highest expression in infected macrophages when compared to other mouse tissues (S11 Fig). Thus, the transcriptional activity of acquired enhancers demonstrated high specificity to the response of BMDM to infection.

**Fig 3.**
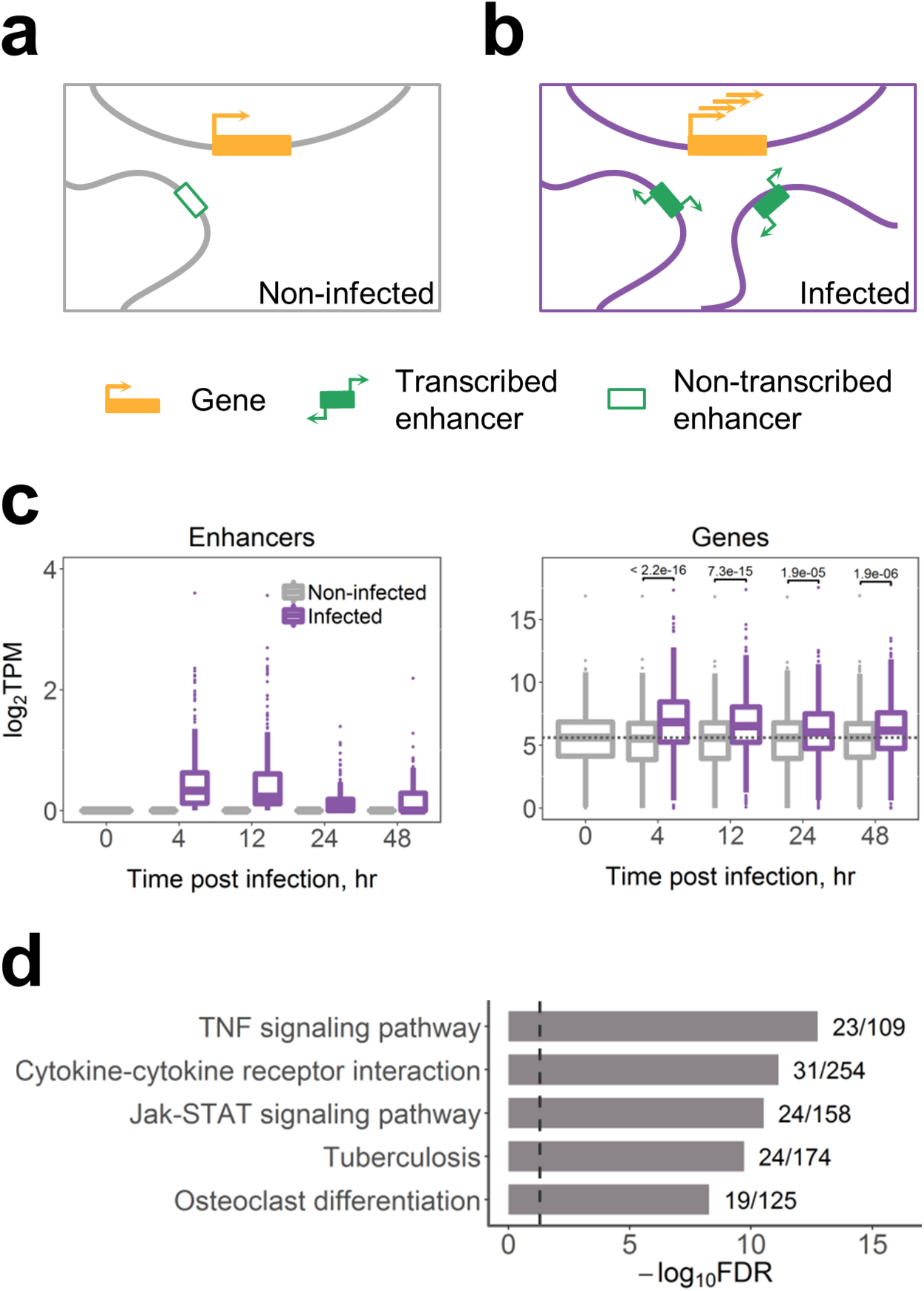
Enhancers that acquire transcriptional activity de novo upon M.tb infection. (a) and (b) show presumable changes in gene regulation upon infection: (a) In non-infected macrophages, a transcriptionally inactive enhancer loops towards its target gene, (b) Upon M.tb infection, the enhancer acquires transcriptional activity; an additional loop is formed de novo for another acquired transcribed enhancer; the gene expression is induced. (c) eRNA expression of 356 acquired enhancers (left) and their 526 target genes (right); dashed line shows median gene expression prior to the infection, expression in TPM was averaged across replicates, p-values of Wilcoxon two-sided rank sum tests are shown. (d) Top 5 KEGG pathway maps with the lowest FDR enriched for 526 target genes of the acquired enhancers; next to the bars are the numbers of genes in the KEGG term covered by our gene list; dashed line indicates FDR = 0.05.

We further compared acquired enhancers to genomic regions carrying H3K4me1 and H3K27ac histone marks, which demarcate pre-established enhancer regions and active enhancers, respectively. We used data from Ostuni et al. [61] for untreated and LPS-treated macrophages. Of 356 acquired enhancers, 83.1% and 99.2% overlapped H3K4me1-enriched regions in untreated and LPS-treated macrophages, respectively, indicating that most acquired transcribed enhancers might be established in naïve macrophages, prior to infection. Unexpectedly, as much as 63.8% of acquired enhancers overlapped H3K27ac-enriched regions in untreated macrophages. However, this percentage was higher at 86% in LPS-treated macrophages, and the corresponding H3K27ac ChIP-seq peaks were stronger enriched in LPS-treated as compared to untreated macrophages (S12 Fig).

### Acquired enhancers in the regulation of immune genes during *M.tb* infection

The acquired enhancers in infected macrophages were associated with 526 genes. The associated genes showed an overall increased expression upon *M.tb* infection (Fig 3c, right panel) and, importantly, a strong enrichment for immune response-related functions (Fig 3d). For further analyses, we sub selected target DEGs that showed up-regulation at 4 h post infection (251 genes, 47.7%, S7 Table).

First, we investigated enhancer-gene associations and found that, at maximum, a DEG was associated with six acquired enhancers. We identified five such genes (*Hivep1*, *Itgb8*, *Pla2g4a*, *Ptgs2*, and *Tnfaip3*). Among the genes, *Pla2g4a* and *Ptgs2* were co-regulated by the same set of acquired enhancers within a TAD (S13 Fig). Both genes are known to be involved in arachidonic acid metabolism, one of the regulators of cell death, and to play a role in infection responses [62]. While *Pla2g4a* showed a moderate induction of log_2_FC = 2.9, expression of *Ptgs2* was induced dramatically with log_2_FC = 11.5 at 4 h post infection (S13a Fig), hinting at its importance during infection.

The strongest induction of log_2_FC = 12.3 at 4 h was observed for endothelin (*Edn1*), a DEG associated with five acquired enhancers (S14 Fig). Edn1 is a well-known vascular regulator; however, its particular roles in infectious diseases including tuberculosis are only beginning to be elucidated [63]. *Edn1* is co-regulated with DEG *Hivep1*, a transcriptional regulator for which the precise function in infected macrophages is unknown (S14 Fig).

All of *Pla2g4a*, *Ptgs2*, *Edn1*, and *Hivep1* genes were additionally associated with other enhancers, which were not classified as acquired enhancers. Among those, *Edn1* and *Hivep1* were associated with one enhancer that was deemed induced in our study (S14c Fig), while *Pla2g4a* and *Ptgs2* were associated with four such induced enhancers (see S13c Fig for eRNA expression of one of them). These enhancers, in contrast to the acquired ones, showed nonzero (although very low) eRNA expression in non-infected macrophages. Notably, in infected macrophages these induced enhancers had a higher expression than the acquired enhancers associated to the same genes (S13-S14 Figs). Thus, up-regulation of DEGs *Pla2g4a*, *Ptgs2*, *Edn1*, and *Hivep1* could not be attributed exclusively to the activity of the acquired enhancers.

We further asked whether any of the 251 up-regulated DEGs were associated exclusively with acquired enhancers. We identified 22 such genes regulated by a total of 18 acquired enhancers. However, in most cases, we observed either low or inconsistent eRNA expression among replicates. Hence, our data could not reliably infer up-regulated DEGs driven exclusively by acquired enhancers. Moreover, the 251 DEGs were associated on average with 1.6 acquired enhancers and 6.1 other enhancers, not classified as acquired. These findings suggest that upon *M.tb* infection, *de novo* transcription at enhancers targeting up-regulated DEGs is acquired in addition to already established transcriptionally active enhancers.

TF binding motif analysis of the acquired enhancers showed overall similar results to that of the induced enhancers, except for Irf1 motif which was over-represented only in induced enhancers, and three TF motifs over-represented only in acquired ones. Among these, a motif for Stat3, a TF known to be involved in *M.tb* infection response [64], overlaps 36.2% of the acquired enhancers. Macrophage-restricted TF Tfec with an overlap of 35.7% has been reported as an important regulator of IL-4 inducible genes in macrophages but was also up-regulated in response to LPS treatment [65]. Finally, the Srebf2 motif overlaps 25.3% of the acquired enhancers. Interestingly, this TF is a host gene of miR-33, a miRNA induced in macrophages by *M.tb* to inhibit pathways of autophagy, lysosomal function and fatty acid oxidation to support *M.tb* intracellular survival [66]. Taken together, these results uncover a novel role of these TFs in the response to *M.tb* infection in BMDM.

## Discussion

Studies in multiple cell types unravelled the fundamental importance of enhancer regions as DNA regulatory elements, however, our current understanding of these elements remains incomplete. High tissue specificity of enhancers is a major hurdle towards establishing a comprehensive catalogue of the full enhancer population [9, 10]. Moreover, emerging evidence indicates that enhancers selectively act in a stimuli- or condition-specific manner [19, 32]. Enhancers often mediate cell-type-specific processes [32]. Previously we reported on the role of transcribed enhancers in macrophage activation and polarisation towards pro- and anti-inflammatory phenotypes [33]. Another recent study linked a specific class of enhancers to the immune response in human [67]. Hence, we hypothesised that enhancers might also regulate the macrophage response to the infection with intracellular pathogens such as *M.tb*. To investigate this possibility, here we analysed *M.tb*-induced changes of gene expression and enhancer activity in macrophages. Our results suggest that transcribed enhancers have a strong influence in the infection response and mediate up-regulation of many important immune protein-coding genes. The strongest macrophage response to *M.tb* was observed at 4 h post infection, hence, we elected to focus on DEGs up-regulated at this time point and to analyse their associated enhancers. We characterised highly transcriptionally induced enhancers and showed that many genes acquired *de novo* transcribed enhancers upon *M.tb* infection. We reported enhancers targeting known immune genes crucial for the genetic response of the host to *M.tb* and highlighted transcription factors that are likely regulating these enhancers. These findings were extended by highlighting particular chromosomal domains carrying groups of highly transcriptionally induced enhancers and genes with previously unappreciated roles in *M.tb* infection.

Previously we have demonstrated that regulation by many enhancers was a concomitant of higher gene expression and tissue-specific functions [33], in agreement with a model of additive enhancer action [8, 68]. Unexpectedly, here we report a similar observation for a highly function-specific set of DEGs up-regulated upon *M.tb* infection. Furthermore, our results indicate that activation of SEs might have a prominent role in regulating macrophage responses to the pathogen, in line with current views of SEs as genomic regions of extreme importance for the regulation of key genes involved in cell-specific processes and responses [35, 36].

Several studies have reported on enhancers that were activated *de novo* upon stimuli [61, 69]. These might represent a particularly functionally important class of enhancers responsible for establishing stimuli-specific gene expression programmes. Ostuni et al. [61] uncovered a set of latent enhancers that lacked any enhancer characteristics in naïve mouse macrophages, but gained active enhancer marks in response to stimulation. Similarly, Kaikkonen et al. [69] identified enhancers activated *de novo* in mouse macrophages stimulated with TLR4 agonist and, interestingly, suggested that eRNA transcription might precede H3K4me1 deposition. In this study, we asked whether any enhancers were non-transcribed in naïve macrophages and acquired *de novo* eRNA transcription upon *M.tb* infection. Interestingly, in contrast to Ostuni et al. [61] and Kaikkonen et al. [69], we found that most of the acquired enhancers might be already marked with H3K4me1 (hence, primed) in naïve macrophages. The remaining 60 of 356 enhancers might acquire both, a H3K4me1 mark and transcriptional activity, upon infection. In agreement with this idea, all 60 enhancers carried H3K4me1 histone marks in LPS-treated macrophages. Moreover, we found that 63.8% of acquired enhancers overlap H3K27ac histone marks in untreated macrophages. This is an unexpectedly large percentage, since H3K27ac is believed to demarcate active enhancers. One possible explanation is that H3K27ac-marked enhancers might have a spectrum of activation states, including those with and without eRNA production. In agreement with this hypothesis, we observe a much stronger H3K27ac enrichment in regions overlapping acquired enhancers in LPS-treated as compared to untreated macrophages. Hence, the strength of H3K27ac enrichment rather than the presence or absence of this histone mark could demarcate actively transcribed enhancers.

Our findings indicate that up-regulated genes in *M.tb*-infected macrophages might acquire *de novo* transcribed enhancers in addition to already established actively transcribed enhancers. We hypothesise that acquired enhancers might be involved in regulating their target genes via at least two different mechanisms. First, activation of acquired enhancers might involve considerable rearrangement of chromatin to allow formation of novel DNA loops between enhancers and their target promoters. Indeed, examples of stimuli-driven dynamical changes in chromatin conformation in mouse macrophages were reported recently [70]. The second hypothetical mechanism would involve the transcriptional activation of enhancers within pre-established chromatin loops. We found that acquired enhancers are often surrounded by other enhancers that are transcribed in naïve macrophages, including *M.tb*-induced enhancers. The fact that these enhancers, at least in some cases, are located close to each other and within SEs points to a hypothetical regulatory mechanism that involves an expansion of active enhancer regions. For instance, a few individual enhancers within a SE might be primed and generate low levels of eRNAs in naïve macrophages. Upon *M.tb* infection, these individual enhancers could serve as ‘seeds’ to enable broader neighbouring regions to acquire enhancer histone marks and stronger eRNA transcription. Such a phenomenon has been described in mouse stem cells, where seed enhancers were shown to expand into SEs [71]. Similarly, a seed enhancer required for activation of a SE has been reported in mammary glands [72]. However, the associated mechanisms and abundances of such seed enhancers remain to be elucidated.

We separately considered two overlapping subsets of enhancers: acquired and induced enhancers. The identification was based on eRNA expression levels before and after *M.tb* infection. However, it is important to note that there is a narrow margin separating these classes, which is influenced by the limits of expression versus noise detection by CAGE and by our sample composition. In other settings, the composition of these classes might differ from our results. For instance, some induced enhancers showed very low (close to zero) eRNA expression in non-infected macrophages, which could be, alternatively, attributed to transcriptional noise.

Signalling pathways regulating macrophage responses to infection have been extensively studied [1, 5, 73], and here we report *M.tb*-induced enhancers that might activate these pathways. We find that induced enhancers might extensively control Tnf and NF-κB signalling pathways by targeting their components, starting from receptors (*Cd14* and *Cd40*) and ligands (*Il1b*, *Tnfrsf1b*, *Tnf*), through mediators (*Traf5*, *Irak2*, *Jak2*, *Malt1*, *Ripk2*, and *Src*), ending with TFs (*Nfkb1* and *Rela*) and numerous pathway effectors. These pathways are known to be activated upon macrophage recognition of *M.tb* and play central roles in shaping immune responses, as they mediate production of pro-inflammatory cytokines and chemokines, and regulate apoptosis [74, 75]. Interestingly, induced enhancers might also control negative feedback regulators of these pathways (*Nfkbia*, *Tnfaip3*, and *Socs3*), which might implicate induced enhancers in terminating immune responses.

As important examples, we highlighted genes regulated by multiple induced or acquired enhancers. We also reported on TADs, where induced enhancers were over-represented, as these chromosomal regions could be considerably affected by *M.tb*. Notably, in this manner we highlighted a group of genes that might be decisive in *M.tb* death versus survival balance via different mechanisms. Knowledge on the regulation of these genes is extremely important for understanding *M.tb* survival strategies and development of novel treatments. Genes with known immune functions are often co-regulated with DEGs with previously unappreciated functions in *M.tb* infection response (such as *Fbxl3*, *Tapt1*, *Edn1*, and *Hivep1*), and these DEGs are, thus, good candidates for further functional studies.

*M.tb* is known to control macrophage cell death pathways, and existing evidence suggests that *M.tb* might induce necroptosis, which facilitates the spread of the pathogen [76]. Here, we found that induced enhancers might be involved in modulating macrophage cell death. For instance, *Tnf* is targeted by three induced enhancers, and might activate both apoptosis and necroptosis via Tnf-signalling pathway, depending on expression of other factors [76]. Activation of a DEG *Tnfrsf1b*, associated with eight induced enhancers, is known to interfere with apoptosis and sensitise macrophages for Tnfr1-mediated necroptosis [37, 38]. In addition, *Pla2g4a*, targeted by four induced enhancers, is involved in metabolism of arachidonic acid, a precursor of lipoxins, leukotrienes, and prostaglandins, lipid mediators which regulate apoptotic/necroptotic balance [62, 77]. *Il1a* and *Il1b* DEGs, co-regulated by four induced enhancers, stimulate production of prostaglandins, linked to necroptosis suppression [77].

Finally, we investigated the transcriptional regulation of induced and acquired enhancers. We identified TFs with binding sites significantly over-represented in these enhancer sets. Importantly, most of these TFs are known to be activated in response to infection, for instance, via Toll-like and Nod-like receptors upon recognition of the pathogen. These findings propose a mechanistic link between *M.tb* infection and transcriptional activation of enhancers that mediate up-regulation of immune genes. Interestingly, we found that most of the TFBS motifs over-represented in induced enhancers were also over-represented in promoters of their target genes, indicating co-regulation of enhancer and promoter transcription by the same cellular machinery.

Macrophages are versatile immune cells, and a spectrum of their phenotypes has been observed, including distinct populations of tissue resident macrophages [78]. *In vivo,* host alveolar macrophages, which are functionally different from BMDM, are infected by *M.tb*. While alveolar macrophages can be easily isolated from mice, the yield is low for a full-scale transcriptomic analysis. In addition, alveolar macrophage transcriptomics is severely affected by the sanitary conditions of the animal facility. Therefore, here we elected to use BMDM as primary macrophages. BMDM are not dependent on health condition of the donor mice, and this choice enabled us to obtain high numbers of cells required for the CAGE analysis. These advantages have also been appealing to other researchers, and BMDM have been used as the primary macrophage model in many immunological transcriptomic studies [79-81]. However, as a consequence of using naïve BMDM as a model, responses observed in our data might differ from host alveolar macrophage responses. Furthermore, some of the transcriptomic changes analysed here could be triggered not by the contact with *M.tb* per se, but rather by other *M.tb* response-associated events, such as cytokine secretion. Future studies of *M.tb* infection in combination with cytokine stimulation could help to further characterise this.

One of the crucial areas of TB research is the development of novel strategies for host-directed therapies, which can stimulate host antimicrobial pathways and suppress host subversion by *M.tb* [82, 83]. Targeting disease-specific enhancers has been investigated as a therapeutic approach in cancer and autoimmune diseases [84, 85]. This study suggests that both acquired and induced enhancers regulate immune genes, which are crucial for *M.tb* survival versus elimination balance. Moreover, transcriptional activity of these enhancers is characterised by a high macrophage- and infection-specificity. Hence, these enhancers are likely good candidate regulatory genomic regions for targeted manipulation of macrophage responses to *M.tb* infection.

## Conclusions

*M.tb* triggers extensive changes in macrophage gene expression programmes that are decisive for the infection outcome, yet the associated regulatory mechanisms remain largely unknown. This is the first to our knowledge study of the role of transcribed enhancers in macrophage response to *M.tb* infection. It extends current understanding of the regulation of *M.tb* responses by linking *M.tb*-responsive transcription factors to activation of transcribed enhancers, which, in turn, target protein-coding immune genes upon infection. Given the increasing promise for enhancer- and chromatin-directed therapy, this work paves the way for further targeted studies towards a host-directed therapy and novel tuberculosis treatments.

## Methods

### Bone marrow-derived macrophage (BMDM) generation

BALB/c mice were purchased from Jackson Laboratories and bred at the Research Animal Facility, University of Cape Town, South Africa. BMDM were generated from 8-12 week old male BALB/c mice as described previously [86].

### Ethics Statement

Mice were sacrificed in accordance with the Animal Research Ethics of South African National Standard (SANS 10386:2008) and University of Cape Town of practice for laboratory animal procedures. The protocol (Permit Number: 012/036) was approved by the Animal Ethics Committee, Faculty of Health Sciences, University of Cape Town, Cape Town, South Africa.

### *M.tb* infection

BMDM were plated in 6-well plates (Nunc, Denmark) at 2 x 10^6^ cells per well and left to adhere for 40 hours. BMDM were then infected with log phase *M.tb* HN878 (MOI = 5) for 4 hours. Cells were washed to remove extracellular mycobacteria and replenished with fresh medium containing 10 μg/ml of gentamycin. At 0, 4, 12, 24 and 48 hours, *M.tb*-infected and non-infected BMDM were lysed with 700 μl of Qiazol (Qiagen, Valencia, CA, USA) for RNA extraction. Total RNA was prepared using miRNAeasy kit (Qiagen, Valencia, CA, USA) and concentration and quality of each RNA samples was verified as described previously [86]. All *M.tb* infection experiments were performed at the Biosafety Level 3 (BSL-3) laboratory, Institute of Infectious Disease and Molecular Medicine (IDM), University of Cape Town, South Africa.

### Data

Macrophage samples were profiled by us using cap analysis of gene expression (CAGE) as described in Roy et al. [87]. Samples used in this study include three biological replicates per time point profiled at 4, 12, 24, and 48 hours post infection in *M.tb* HN878-infected and control macrophages (except for 48 h infected samples, where two biological replicates were available). In addition, four biological replicates were profiled prior to infection at 0 h and six more samples were profiled during macrophage cultivation before this time point.

Mouse genome assembly mm10 and Ensembl gene models version 75 were used [88]. CAGE-derived tag counts were normalized to tags per million (TPM) using TMM normalization [89].

Data were processed, including identification of enhancer regions and enhancer-gene associations, as described in Denisenko et al. [33]. Briefly, enhancers were defined following the strategy of Andersson et al. [17] as bidirectionally transcribed 401 bp regions, and further were required to overlap ChIP-seq-derived H3K4me1 histone marks [61]. Enhancer-gene associations were established by selecting enhancers and promoters which were located within the same TAD [28] and showed positive Spearman’s correlation coefficient of expression in macrophages with FDR < 10^-4^ (Benjamini-Hochberg procedure [90]). Of all enhancer-gene associations established in [33], we here sub selected only those with a positive Spearman’s correlation of expression specifically in the infected macrophage samples.

### Differential expression analysis

Differential gene expression analyses were performed using the exact test implemented in edgeR [89]. Four macrophage samples profiled prior to the infection (0 h) were used as a control. The p-values were adjusted for multiple hypothesis testing using the Benjamini-Hochberg procedure [90]. FDR ≤ 0.05 and log_2_ fold change > 1 (< −1) thresholds were used to select differentially expressed up-(down-) regulated genes (DEGs).

### Gene set enrichment analysis (GSEA)

KEGG pathway maps [91] were used as a set of biological terms for GSEA. We used the hypergeometric distribution to calculate the probability of obtaining the same or larger overlap between a gene set of interest and each biological term [92]. Derived p-values were corrected for multiple testing using Benjamini-Hochberg procedure [90]. As a background gene list, a set of 22,543 Ensembl protein-coding genes (version 75) was used [88].

### Overlaps with ChIP-seq data

We used ChIP-seq data for H3K4me1 and H3K27ac histone marks profiled in untreated and LPS-treated macrophages by Ostuni et al. [61] (Gene Expression Omnibus accession GSE38379). Genomic coordinates of significant ChIP-seq peaks were converted from mm9 to mm10 using the liftOver program [93].

### Transcription factor binding analysis

Transcription factor (TF) binding profiles were downloaded from JASPAR database, 7th release, 2018 [94]. The Clover program [95] was used for identification of statistically over-represented motifs. Enhancer regions were tested against three background DNA sets, as previously defined by us [33]: 1) the whole set of transcribed mouse enhancers; 2) a subset of these enhancers not transcribed in macrophages; 3) a set of random genomic regions excluding gaps, repeated sequences, Ensembl coding regions, and the transcribed mouse enhancers. Promoter regions were tested against the following three sets: 1) all promoters expressed in mouse tissues; 2) a subset of those not expressed in macrophages; 3) the same set of random genomic regions as used for enhancers. Promoters were used as defined in [33] and were extended by 500 bp upstream and downstream. Motifs with p-value < 0.01 for each of the three background sets were selected as significantly over-represented. TFs that were significantly differentially expressed and up-regulated at 4 h post infection when compared to 0 h were retained.

### *M.tb*-induced and acquired enhancers

*M.tb*-induced enhancers were selected among those associated with DEGs up-regulated at 4 h post infection. Mean eRNA expression for these enhancers at 4 h and its fold change compared to 0 h were calculated. Enhancers were defined as induced, if both these values were in the upper quartiles of their corresponding distributions. Acquired enhancers were defined as those with no detectable eRNA expression in each of 22 non-infected BMDM samples, and nonzero expression in any of the infected macrophage samples.

### TADs enriched for enhancers

Genomic coordinates of TADs in mouse embryonic stem cells were obtained from a study by Dixon et al. [28] and were converted from mm9 to mm10 using the liftOver program [93]. To uncover chromosomal domains that might be important in macrophage response to *M.tb*, we identified TADs that were significantly enriched for induced enhancers. A hypergeometric test was performed for each TAD by comparing the total number of BMDM enhancers in that TAD to the subset of those deemed induced. The p-values for 1,228 TADs were corrected for multiple hypothesis testing using Benjamini-Hochberg procedure [90]. TADs with FDR < 0.05 were selected as significantly enriched for induced enhancers.

## Declarations

### Consent for publication

Not applicable.

### Availability of data and materials

The dataset analysed in the study is available in the FANTOM5 repository, http://fantom.gsc.riken.jp/5/datafiles/reprocessed/mm10_v2/basic/. The datasets supporting the conclusions of this article are included within the article and its additional files.

### Authors’ contributions

ED performed computational analyses. SS designed the study. RG and HS performed the experiments. SS and ED analysed data, interpreted results, and wrote the manuscript with input from all authors. RG, MM, HS, and FB helped interpret results and provided data. All authors read and approved the final manuscript.

## Acknowledgements

Not applicable.

## Supporting information

S1 Fig (.tif). Many enhancers respond to *M.tb* infection with increased eRNA expression.

S2 Fig (.tif). Higher number of associated enhancers is a concomitant of higher gene expression and immune functions in infected macrophages.

S3 Fig (.tif). Up-regulated DEGs associated with super enhancers show more infection-specific functions.

S4 Fig (.tif). 257 induced enhancers associated with 263 DEGs up-regulated at 4 h post infection.

S5 Fig (.tif). Expression of the induced enhancers in mouse tissues. S6 Fig (.tif). Regulation of Irg1, Cln5, and Fbxl3 genes.

S7 Fig (.tif). Regulation of Hilpda gene. S8 Fig (.tif). Regulation of Itgb8 gene.

S9 Fig (.tif). Regulation of Cd38, Bst1, and Tapt1 genes.

S10 Fig (.tif). Regulation of Ccl9, Ccl3, Ccl4, and Wfdc17 genes.

S11 Fig (.tif). Expression of the acquired enhancers in mouse tissues. S12 Fig (.tif). H3K27ac ChIP-seq peaks.

S13 Fig (.tif). Regulation of Pla2g4a and Ptgs2 genes. S14 Fig (.tif). Regulation of Edn1 and Hivep1 genes.

S1 Table (.xlsx). A list of DEGs up-regulated at any time points, with their associated enhancers in infected macrophages.

S2 Table (.xlsx). KEGG pathway maps significantly enriched for up-regulated DEGs associated with more than two transcribed enhancers.

S3 Table (.xlsx). Induced enhancers with associated target DEGs up-regulated at 4 h post infection.

**S4 Table (.xlsx). A full list of non-macrophage mouse samples split by tissue.** Tissues with at least ten samples were considered separately, the rest of the samples were combined together into an ‘Others’ category.

S5 Table (.xlsx). TADs enriched for induced enhancers.

**S6 Table (.xlsx). Three selected KEGG pathway maps enriched for DEGs regulated by induced enhancers.** Corresponding DEGs and induced enhancers are listed along with correlation coefficient and p-value.

S7 Table (.xlsx). Acquired enhancers with associated target genes up-regulated at 4 h post infection.

